# Neuromodulatory silencing of nerve terminals

**DOI:** 10.1101/2022.09.16.508300

**Authors:** Daniel C. Cook, Timothy A. Ryan

## Abstract

Control of neurotransmission efficacy is central to theories of how the brain computes and stores information. Neuromodulators are critical in this problem as they locally influence synaptic strength and can operate on a wide range of time scales. Presynaptic function is heavily influenced by G-protein coupled receptors (GPCRs) that in part restrict voltage-gated calcium (Ca^2+^) influx in the active zone. Here, using quantitative analysis of both single bouton Ca^2+^ influx and exocytosis, we uncovered an unexpected non-linear relationship between the magnitude of action potential driven Ca^2+^ influx and the concentration of external Ca^2+^ ([Ca^2+^]_e_). We find that this unexpected relationship is leveraged by neuromodulator signaling when operating at the nominal physiological set point for [Ca^2+^]_e_, 1.2 mM, to achieve complete silencing of nerve terminals. These data imply that the information throughput in neural circuits can be readily modulated in an all-or none fashion at the single synapse level when operating at the physiological set point.

## Introduction

Neuromodulators serve a key control function in the brain, acting through synaptic GPCRs to modify the efficacy or time course of neurotransmission through pre- and postsynaptic mechanisms^1^. Neuromodulators are often secreted near synapses and exert their effects locally^2–4^. They can thereby modify region-specific synaptic properties and change the weighting of targeted synapses, in turn altering local computation. A canonical mode of presynaptic modulation occurs via the binding of voltage-gated calcium channels (VGCCs) by G-protein βγ subunits after they dissociate from the trimeric G-protein complex following GPCR activation^5^. The impact of GPCR-mediated inhibition of VGCCs on neurotransmission^6,7^ is amplified by the exquisite sensitivity of exocytosis to the magnitude of Ca^2+^ influx, which underlies the ability to rapidly change exocytotic rates at sites of neurotransmitter release. This sensitivity is driven by the cooperative binding of multiple Ca^2+^ ions to one or more Ca^2+^ sensors that form core elements of the exocytic machinery^8–11^. Modifying the open-probability of VGCCs therefore serves as a potent means of presynaptic control, as small changes in Ca^2+^ influx can lead to large changes in neurotransmitter release.

The impact of Ca^2+^ on exocytosis depends on local changes in Ca^2+^ concentration near the active zone, which in turn depend on the external Ca^2+^ concentration bathing the nerve terminal^12^. Thus, the steepness of the relationship between exocytosis rates and Ca^2+^ influx depends crucially on resting [Ca^2+^]_e_. In vivo, plasma [Ca^2+^] is strictly regulated, and the CNS is very sensitive to plasma [Ca^2+^] perturbations, with seizures precipitated by hypocalcemia and psychiatric symptoms progressing to lethargy or coma with worsening hypercalcemia^13^. Feedback loops based on signaling Ca^2+^ sensors control of the parathyroid gland act to maintain plasma [Ca^2+^] at a set point of 1.2 mM in most mammals^14^. Here we used quantitative optical tools to investigate the sensitivity of exocytosis rates at the single-synapse level when operating at the physiological set point for [Ca^2+^]_e_. In a narrow range of [Ca^2+^]_e_ near the physiologic concentration, our experiments revealed an unexpected sub-proportionality of Ca^2+^ influx relative to [Ca^2+^]_e_, indicating that a minimal Ca^2+^ entry is needed to sustain robust Ca^2+^ influx during AP firing. We show that three different approaches to modestly decrease Ca^2+^ influx, 1) blocking a fraction of Ca^2+^ channels, 2) lowering [Ca^2+^]_e_, or 3) application of neuromodulators known to reduce Ca^2+^ influx leads to complete silencing of a subset of nerve terminals. Thus, in this operating regime, neuromodulators that normally decrease presynaptic Ca^2+^ current now act as digital switches to eliminate single bouton synaptic throughput.

## Results

### Ca^2+^ influx into nerve terminals is not 1:1 proportional to [Ca^2+^]_e_ near the physiologic set point

In order to examine how Ca^2+^ influx driven by action potentials (APs) at nerve terminals is impacted by changes in [Ca^2+^]_e_, we used the genetically-encoded Ca^2+^ indicator physin-GCaMP6f transfected sparsely into primary cultures of dissociated rat hippocampal neurons (Figure 1). We measured the changes in GCaMP6f signal in response to a 20 AP burst stimulus (20 Hz) at 0.8, 1.2 and 2.0 mM [Ca^2+^]_e_ (Figures 1A and B). Signals were corrected for local abundance of the protein by determining the GCaMP6f fluorescence under saturating Ca^2+^ binding (determined following ionomycin application) and converted to absolute intracellular Ca^2+^ concentrations ([Ca^2+^]_i_, see methods). Previous studies investigating the dependency of Δ[Ca^2+^]_i_ to [Ca^2+^]_e_ in nerve terminals of the CNS found the relationship obeyed a simple saturable pore model with a Kd of [Ca^2+^]_e_ ~2.5 mM^11,15^. This model predicts that changes in Δ[Ca^2+^]_i_ will exhibit 1-to-1 proportionality to [Ca^2+^]_e_ at concentrations well below Kd, which is substantially higher than the physiologic set point ([Ca^2+^]_e_ = 1.2 mM). One important confound, however, is that previous measurements were not carried out at physiological temperature. Here, where the temperature was set to 37°C, we observed an unexpected sub-proportional relationship of Δ[Ca^2+^]_i_ to [Ca^2+^]_e_ as the concentration was lowered below 2.0 mM (Figures 1B and C). The magnitude of difference between the expected and experimentally measured Δ[Ca^2+^]_i_ was greater at [Ca^2+^]_e_ 0.8 mM compared to 1.2 mM, highlighting that this discrepancy manifests most prominently below the set point (Figure 1C). A linear extrapolation from of the mean Δ[Ca^2+^] measurements relative to [Ca^2+^]_e_ predicts the cessation of Ca^2+^ entry into nerve terminals at [Ca^2+^]_e_ of 0.47 mM (Figure 1C). Thus, nerve terminals have a non-zero threshold of [Ca^2+^]_e_ below which presynaptic Ca^2+^ influx and function is abolished.

**Figure 1.**
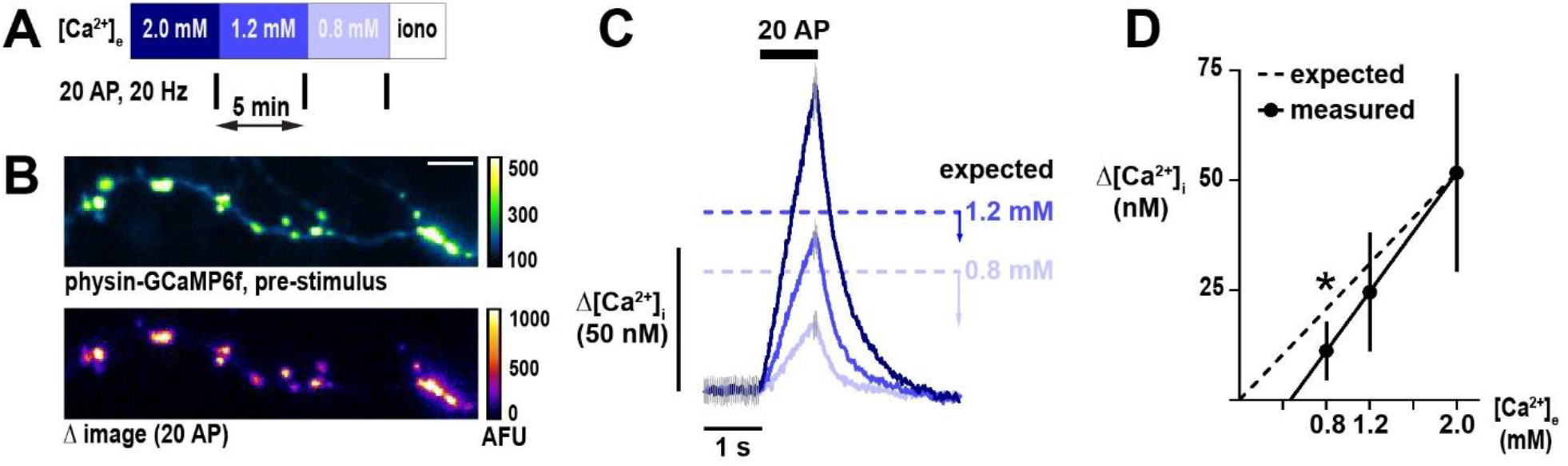
Unexpected sub-proportionality in dependence of presynaptic Ca^2+^ influx on [Ca^2+^]_e_. (A) Schematic of experimental protocol. Neurons are stimulated with a brief AP train (20 AP in 1 s) at 3 different [Ca^2+^]_e_ centered on the physiologic set point of 1.2 mM. Ionomycin, a Ca^2+^ ionophore, is administered at the end of the experiment to convert fluorescence into absolute [Ca^2+^]_i_. (B) An axon from a neuron expressing GCaMP6f localized to nerve terminals by synaptophysin (physin-GCaMP) at rest (top) and a difference image showing the response to 20 AP (bottom). The difference image is the mean peak (5 frames) subtracted by the mean baseline (49 frames) in [Ca^2+^]_e_ 2 mM. For display, a representative subset of terminals was highlighted. Scale bar is 10 μm. (C) Traces of responses to 20 AP from the neuron shown in B, with [Ca^2+^]_e_ color-coded as in A. Expected Δ[Ca^2+^]_i_ at 0.8 and 1.2 mM is represented by dashed lines and calculated as a change in influx relative to 2.0 mM proportional to the ratios of [Ca^2+^]_e_. Arrow bars show the difference between expected and measured Δ[Ca^2+^]_i_. Traces are mean, with error (SEM) represented by gray lines in only pre-stimulus and peak frames for clarity (n = 211 nerve terminals). (D) Summary of Δ[Ca^2+^]_i_ as a function of [Ca^2+^]_e_ (mean ± 95% confidence interval, n = 9 neurons). The x-axis intercept for measured changes in Δ[Ca^2+^]_i_ is 0.47 mM. *p < 0.05, one-sample t-test compared to the expected value.

### Modest changes of [Ca^2+^]_e_ near the physiologic set point controls the proportion of silent nerve terminals

The above measurements represented ensemble averages of all terminals with a measurable signal following saturation of physin-GCaMP. To further probe the apparent non-zero threshold of [Ca^2+^]_e_ that enables Ca^2+^ entry, we assessed physin-GCaMP responses at individual nerve terminals with varying [Ca^2+^]_e_ (Figure 2). Surprisingly, we found that a subset of terminals exhibited an all-or-none response to lowering [Ca^2+^]_e_ below the physiologic set point such that Δ[Ca^2+^]_i_ is selectively abolished (Figure 2B). To rigorously quantify this behavior, we employed a modest cutoff of one standard deviation of the pre-stimulus noise above the mean fluorescence to separate responding terminals from those without measurable synaptic activity, termed silent^16-18^. We observed that silencing occurred despite robust Δ[Ca^2+^]_i_ at higher [Ca^2+^]_e_ and even if synaptic neighbors of a terminal exhibited persistent Ca^2+^ entry at a lower concentration (Figures 2A and B), suggesting the impact of [Ca^2+^]_e_ on neuronal function is exerted at the single synapse level. Applying this approach to a population of neurons (Figure 3), we found substantial heterogeneity in the proportion of silent terminals (coefficient of variation was 38%, 56%, and 57% for [Ca^2+^]_e_ 0.8, 1.2, and 2.0 mM, respectively) but a strikingly disproportionate increase in the mean proportion of silencing below 1.2 mM (Figure 3C). The change in percentage of silencing versus change in [Ca^2+^]_e_ was 48.9% mM^-1^ transitioning from 1.2 to 0.8 mM compared to 19.0% mM^-1^ from 2.0 to 1.2 mM (Figure 3C). As expected, responding terminals demonstrated Δ[Ca^2+^]_i_ that scaled with [Ca^2+^]_e_ while silent terminals were invariant (Figure 3D). Thus, the pronounced, selective silencing of a subset of terminals by lowering [Ca^2+^]_e_ below the physiologic set point suggests that this concentration may be an important lever modulating synaptic function.

**Figure 2.**
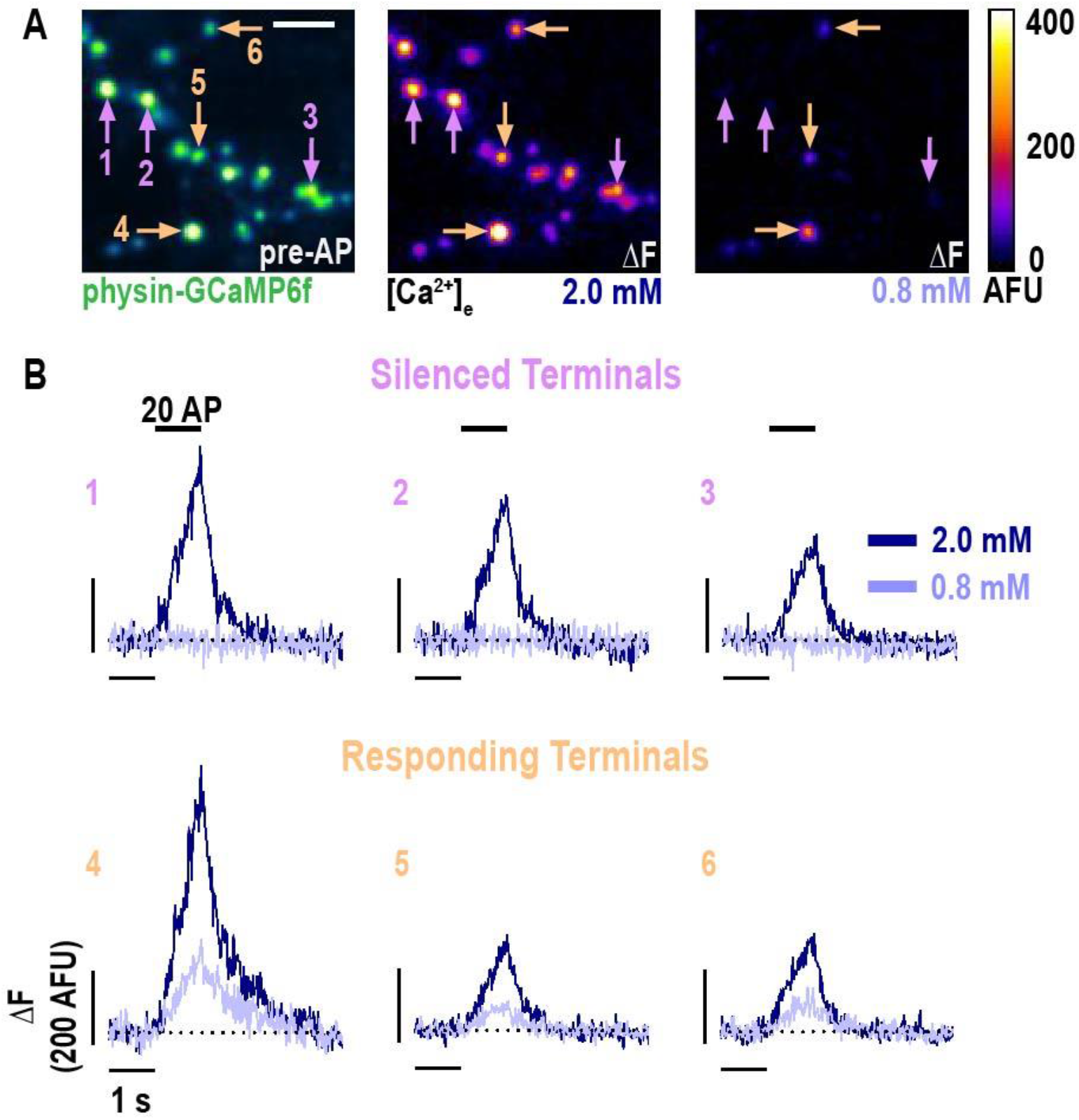
Selective nerve terminal silencing caused by changing [Ca^2+^]_e_ around the physiologic set point. (A) Pre-stimulus average (left) and difference images (right) of AP stimulation (20 APs in 1 s) of a neuron expressing physin-GCaMP. The difference image is the mean peak (5 frames) subtracted by the mean baseline (49 frames) at [Ca^2+^]_e_ 2.0 and 0.8 mM. For display, a subset of representative terminals was selected and a Gaussian convolution with radius of 1 pixel was applied. (B) Traces of the terminals indicated in A, shown as ΔF, demonstrating selective silencing (upper row compared to lower row) of a subset of terminals. Dotted line is pre-stimulus mean fluorescence for each terminal.

**Figure 3.**
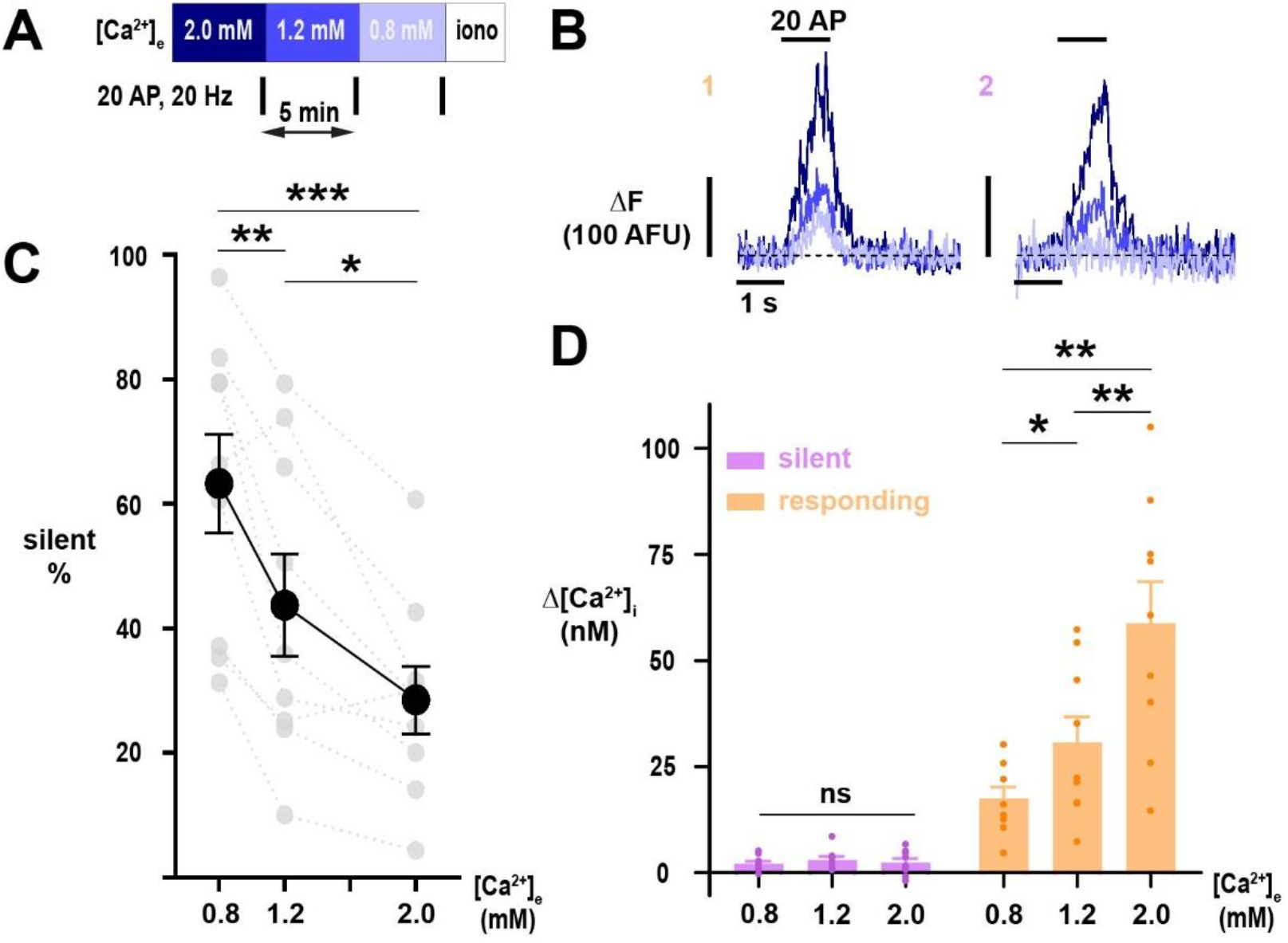
Silencing of Ca^2+^ influx into nerve terminals is potently modulated by changes in [Ca^2+^]_e_ about the physiologic set point. (A) Diagram of experimental protocol. (B) Example traces of single nerve terminals expressing physin-GCaMP, demonstrating persistent responsiveness (1) or silencing (2) with lowering of Ca^2+^ below physiologic extracellular levels. Dotted line is pre-stimulus mean fluorescence (ΔF = 0). (C) The percentage of silent nerve terminals. Black dots are mean, error bars are SEM, and grey dots are individual cells. (D) Summary of Δ[Ca^2+^]_i_, separated into silent or responding terminals. C and D were analyzed with one-way ANOVA and Tukey’s post-test for multiple comparisons, * p < 0.05, ** p < 0.01, *** p < 0.001, n = 9.

### The impact of [Ca^2+^]_e_ on synaptic silencing is exerted through global reduction in Ca^2+^ entry irrespective of VGCC subtype

Because silencing of Ca^2+^ influx occurred in a variable proportion of terminals in each neuron, we next investigated whether this selectivity is mediated by the relative expression of VGCC subtypes. Ca^2+^ influx at hippocampal nerve terminals occurs predominantly via N- and P/Q-type VGCCs, with substantial heterogeneity between neurons in their proportional contribution to Ca^2+^ influx^19–21^. Previously, cyclin-dependent kinase 5 (CDK5) was shown to silence a subpopulation of nerve terminals, an effect mediated by N-type channels only^22^. We therefore hypothesized that N-type channels may exhibit a greater propensity to [Ca^2+^]_e_-driven silencing as compared to P/Q-type channels, accounting for neuronal variability in the proportion of silent terminals. To examine whether [Ca^2+^]_e_-mediated silencing exhibits subtype selectivity, we utilized ω-conotoxin-GVIA and ω-agatoxin IVA, potent toxins that specifically inhibit N- and P/Q-type channels, respectively (Figure 4A). As previously demonstrated, the contribution of presynaptic VGCC subtypes to Δ[Ca^2+^]_i_ is variable between hippocampal neurons (Figures 4B and C)^19^. A subset of terminals is silenced with acute inhibition of either subtype, but isolation of N-type channels leads to greater proportions of silent terminals (Figures 4B and D). To further investigate whether a difference exists between subtypes in the likelihood of silencing, we compared Ca^2+^ entry in responding terminals to the proportion of silent terminals at each tested [Ca^2+^]_e_ (Figures 4C and D). This analysis reveals a non-linear relationship, such that lowering of absolute Ca^2+^ influx steeply increases the fraction of silent terminals. The comparison of Δ[Ca^2+^]_i_ to silencing was well described with a Hill equation (r^2^ = 0.99, Kd = 25 nM, n = −1.79) and closely matched the relationship observed without toxin application (r^2^ = 0.77, Kd = 23 nM, n = −1.46; Figure 4E). This analysis indicates that the proportion of silent nerve terminals is critically related to global absolute Ca^2+^ influx and that differences between N- and P/Q-type channels are driven primarily by the impact that acute inhibition has on Ca^2+^ entry.

**Figure 4.**
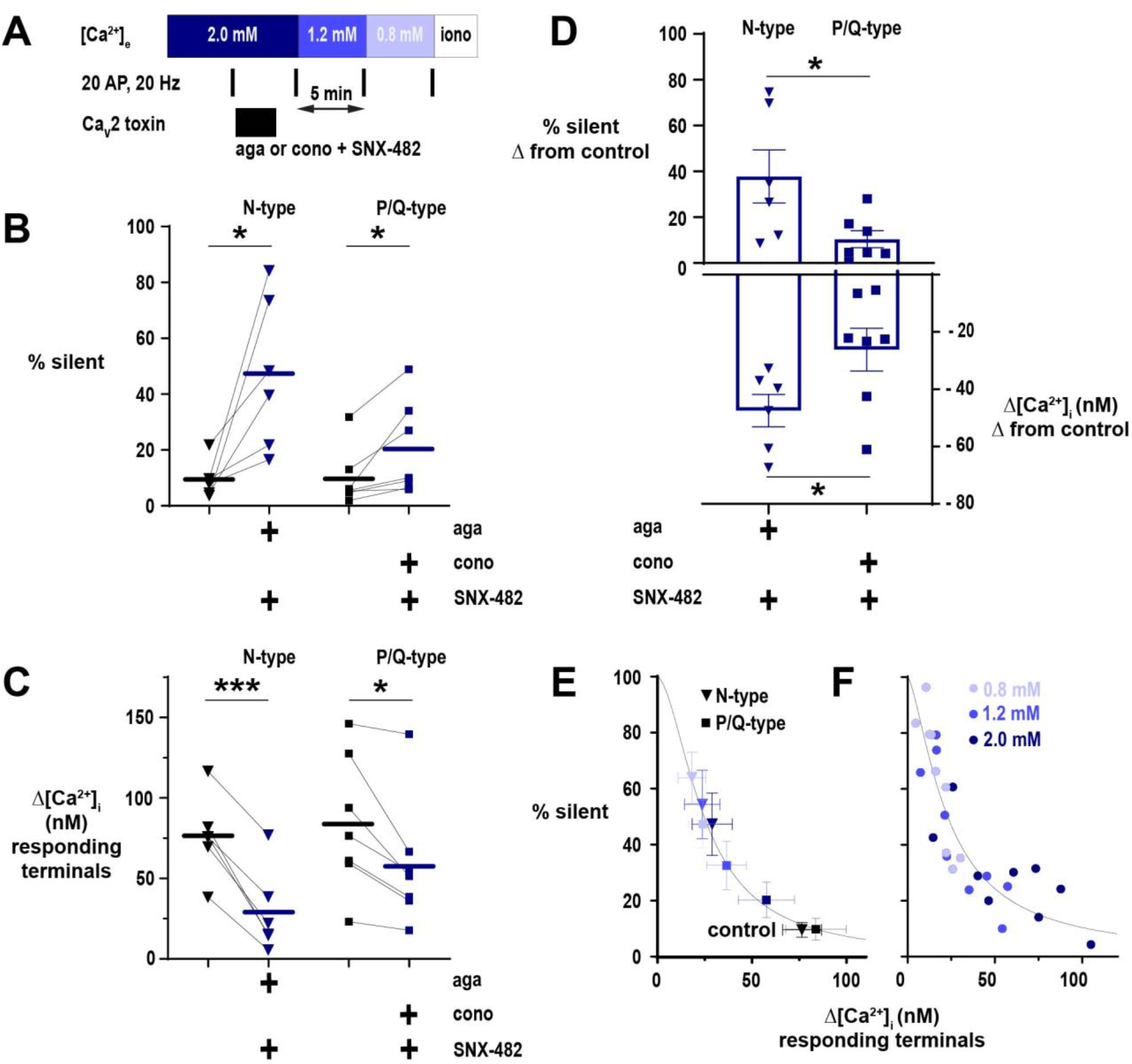
The proportion of nerve terminal silencing in neurons is related to absolute Ca^2+^ influx irrespective of Ca_V_2 subtype. (A) Diagram of experimental protocol applied to neurons expressing physin-GCaMP. (B) The percentage of silent nerve terminals before and after toxin application to isolate a CaV2 subtype at [Ca^2+^]_e_ 2 mM. (C) Summary of Δ[Ca^2+^]_i_ in responding terminals after toxicologic isolation of N-type or P/Q-type channels. (D) Comparison of the effects of toxin application to isolate CaV2 subtypes on silencing (upper graph, shown on left axis) and Ca^2+^ influx (lower graph, shown on right axis). For B-D, symbols are mean of individual neurons, lines and bars are mean of all cells, and error bars are SEM. B and C were analyzed with paired t-test, and D was analyzed with unpaired t-test. * p < 0.05, *** p < 0.001, n = 6 for N-type and n = 7 for P/Q-type. (E) The percentage of silent terminals plotted against Δ[Ca^2+^]_i_ in responding terminals in control (black symbols) and following toxin application at varying [Ca^2+^]_e_. Symbols are mean and error bars are SEM. Data are fit with a Hill model, with the maximum at Δ[Ca^2+^]_i_ = 0 constrained to 100%. (F) Relationship of the percentage of silent terminals to Δ[Ca^2+^]_i_ in responding terminals with varying [Ca^2+^]_e_ in neurons not treated with CaV2 toxins. Dots are mean of individual cells from neurons presented in Figure 2. Data are fit as in D.

### Changing [Ca^2+^]_e_ near the physiologic set point potently controls the silencing of SV recycling and glutamate exocytosis at neurotransmitter release sites

Our observation of steep changes in the proportion of nerve terminals in which Ca^2+^ influx was silenced by lowering [Ca^2+^]_e_ below the physiologic set point led us to investigate the impact of [Ca^2+^]_e_ on neurotransmitter release. We first addressed silencing of SV recycling with pHluorin, a pH-sensitive fluorescent protein^23,24^, tagged to vGLUT1 (vGpH), which is quenched in the acidic lumen of synaptic vesicles but fluoresces with exocytosis and exposure to the relatively alkaline extracellular environment^25^ (Figure 5A-C). We employed a robust stimulus of 200 AP (20 Hz) to distinguish responding from silent terminals^22,26^ (Figures 5A and C). Similar to synaptic Ca^2+^ influx, silencing of SV recycling occurred at a substantial subset of nerve terminals (~25 to 55%, Figure 5D). The proportion of silent terminals was heterogeneous across neurons but the mean and range at each [Ca^2+^]_e_ were similar to the degree of silencing observed with Δ[Ca^2+^]_i_ (Figure 3C). Separating terminals by whether they were silent or responding, the increase in SV recycling scaled with [Ca^2+^]_e_ while silent terminals were invariant (Figure 5E). Thus, as with Ca^2+^ influx, SV recycling is modulated in an all-or-none manner by modest changes in [Ca^2+^]_e_ around the physiological set point.

**Figure 5.**
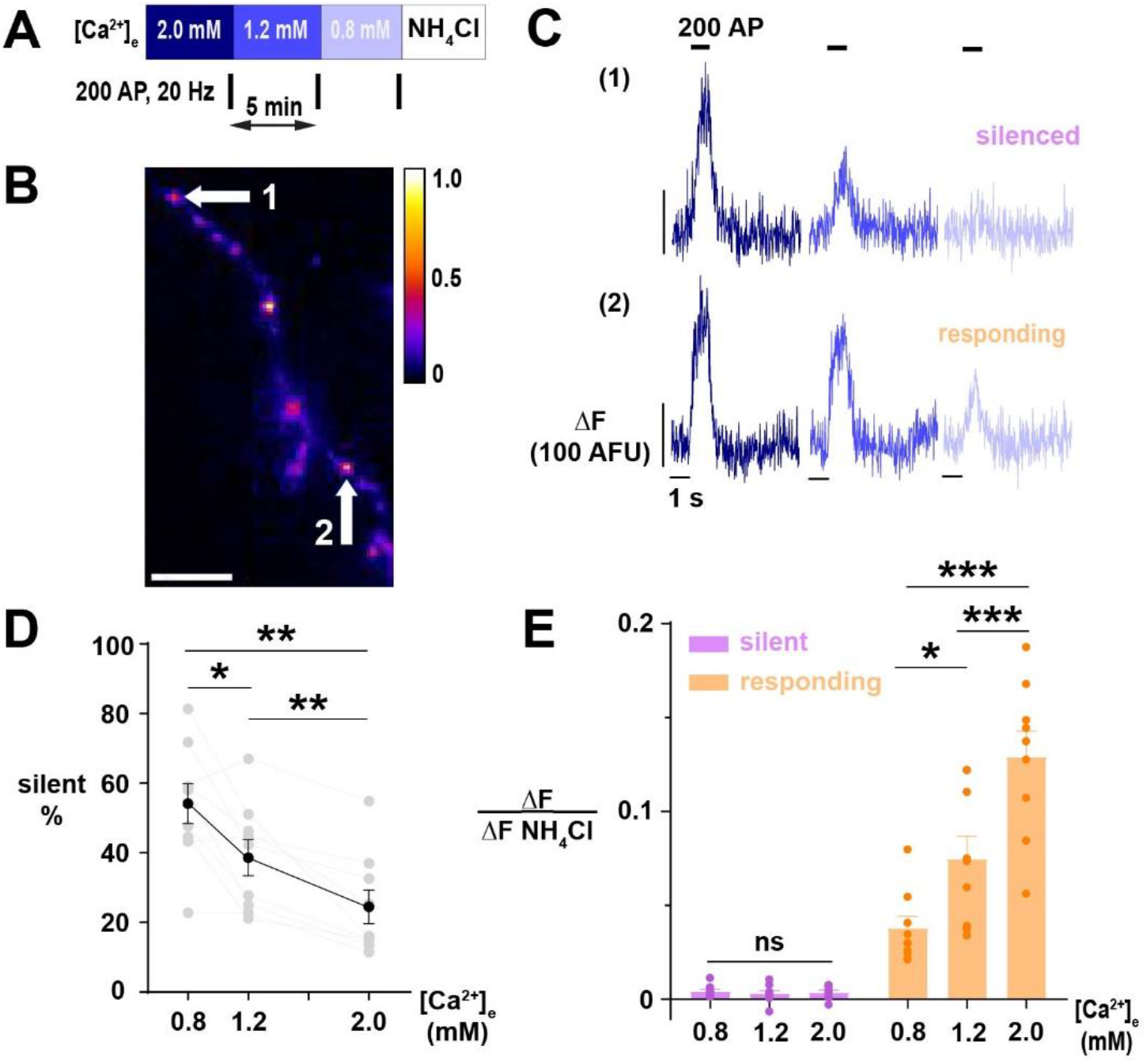
Exocytosis of SVs exhibits [Ca^2+^]_e_-driven silencing. (A) Diagram of experimental protocol. (B) Segment of an axon expressing vGpH, showing difference in fluorescence in nerve terminals revealed by application of NH4Cl 50 mM. Scale bar is 10 μm, and calibration is maximum-normalized fluorescence intensity. (C) SV recycling of individual nerve terminals indicated in B demonstrating selective silencing at [Ca^2+^]_e_ 0.8 mM in (1) compared to persistent response in (2). Traces of different [Ca^2+^]_e_ are color-coded as in A. (D) Percentage of silent nerve terminals. Black dots are mean, error bars are SEM, and grey dots are individual cells. (E) SV exocytosis at different [Ca^2+^]_e_, with terminals separated into responding or silent. Dots are individual neurons, bars are mean, and error bars are SEM. D and E were analyzed with one-way ANOVA and Tukey’s post-test for multiple comparisons, * p < 0.05, ** p < 0.01, *** p < 0.01, ns p > 0.05, n = 9.

We next sought to directly assess how [Ca^2+^]_e_ regulates silencing of glutamate release in the regime of stimulation with a single AP. To do so, we utilized iGluSnFR3 v857 (iGluSnFR), a genetically-encoded, fluorescent biosensor of extracellular glutamate concentrations with an excellent signal-to-noise ratio and temporal resolution enabling quantification of glutamate release from one AP at individual nerve terminals^27^. Co-expression of mRuby-synapsin allowed measurements to be localized to neurotransmitter release sites (Figure 6B); however, because iGluSnFR is anchored by glycophosphatidylinositol (GPI) to the cell surface, the indicator does not necessarily discriminate between glutamate released by the transfected cell or neighboring terminals from non-transfected neurons^27^. Accordingly, while fluorescent signals at mRuby-synapsin puncta may receive contributions from additional terminals, silent terminals are definitively assigned as inactive. With this approach, we found glutamate exocytosis evoked by a single AP exhibited higher proportions of silencing at neurotransmitter release sites as [Ca^2+^]_e_ was lowered, again demonstrating a marked increase in silencing as the concentration decreased below the physiologic set point (Figures 6C and D). Thus, using these two fluorescent biosensors quantifying neurotransmitter handling under different stimulation regimes demonstrates that [Ca^2+^]_e_ is an important regulator of presynaptic function, differentially dictating across synapses and neurons whether neurotransmitter will be released.

**Figure 6.**
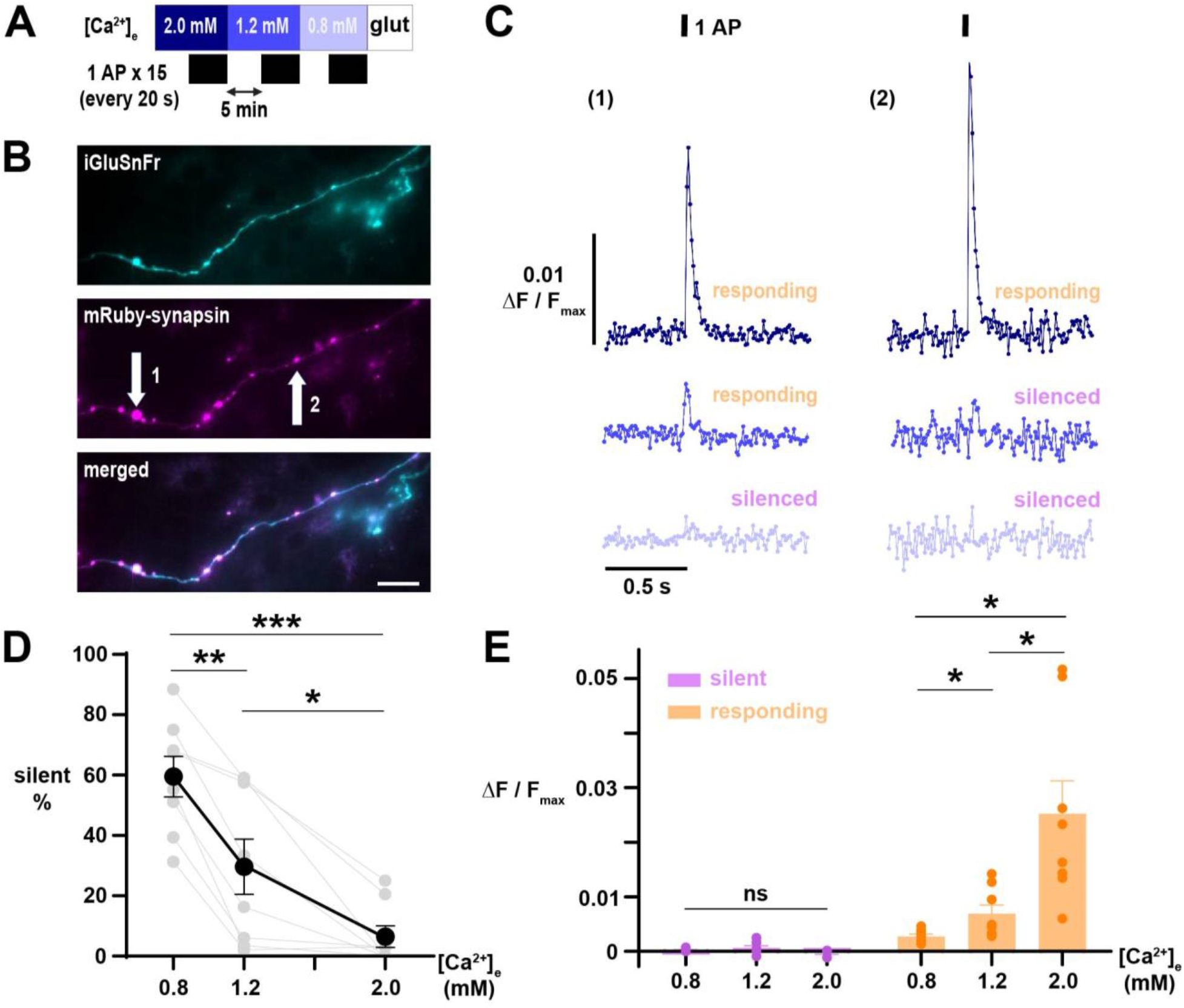
Glutamate release elicited by 1 AP demonstrates [Ca^2+^]_e_-driven silencing at neurotransmitter release sites. (A) Diagram of the experimental protocol. Responses of iGluSnFR-expressing neurons were averaged over 15 single AP trials delivered in ~20 s intervals. (B) Difference image (peak minus baseline fluorescence) of an axon expressing iGluSnFR (cyan, upper) stimulated with 1 AP in [Ca^2+^]_e_ 1.2 mM, with terminals marked by mRuby-synapsin (magenta, middle; merge lower). Scale bar 20 μm. (B) Traces of iGluSnFR measured at nerve terminals indicated in B, illustrating differential silencing as [Ca^2+^]_e_ is lowered. Traces of different [Ca^2+^]_e_ are color-coded as in A. (D) Percentage of silent nerve terminals. Black dots are mean, error bars are SEM, and grey dots are individual cells. (E) Glutamate release, quantified as ΔF/Fmax, as [Ca^2+^]_e_ is varied with terminals separated by responding or silent. Dots are the mean of individual cells, bars are mean, and error bars ± SEM. D and responding terminals in E were analyzed with one-way ANOVA and Tukey’s post-test for multiple comparisons. Silent terminals in E were analyzed with a mixed-effects model because of missing values from neurons without silent terminals. * p < 0.05, ** p < 0.01, *** p < 0.001, ns p > 0.05, n = 8.

### Neuromodulators that lower synaptic Ca^2+^ currents similarly modulate the proportion of silent nerve terminals

The selective silencing of Ca^2+^ entry and neurotransmitter release in subpopulations of nerve terminals by globally reducing Ca^2+^ influx may be an important mechanism determining the behavior of synapses to biological processes affecting Ca^2+^ currents (I_Ca_). For instance, a subset of important neuromodulators affect neurotransmission by lowering synaptic Ca^2+^ influx^28^. One such canonical mechanism is agonism of the GABAB receptor (GABABR), a G-protein coupled receptor (GPCR) that lowers I_Ca_ by approximately 50% via interactions of the Gβγ subunit with presynaptic VGCCs^29–32^. Thus, we hypothesized that GABA_B_R-mediated decreases in I_Ca_ will significantly increase the degree of nerve terminal silencing at the physiologic set point of [Ca^2+^]_e_. We compared the effect of baclofen, a GABABR agonist, on silencing of Ca^2+^ influx and SV recycling using physin-GCaMP and vGpH, respectively (Figures 7A-C). Consistent with our proposed mechanism, agonizing GABA_B_R at [Ca^2+^]_e_ 1.2 mM caused an additional ~20% of terminals to become silent in both Ca^2+^ influx and SV recycling (Figure 7D), similar to the proportional increase we observed decreasing [Ca^2+^]_e_ to 0.8 mM. These results thus confirm that neuromodulators acting presynaptically to decrease I_Ca_ can silence a substantial proportion of nerve terminals.

**Figure 7.**
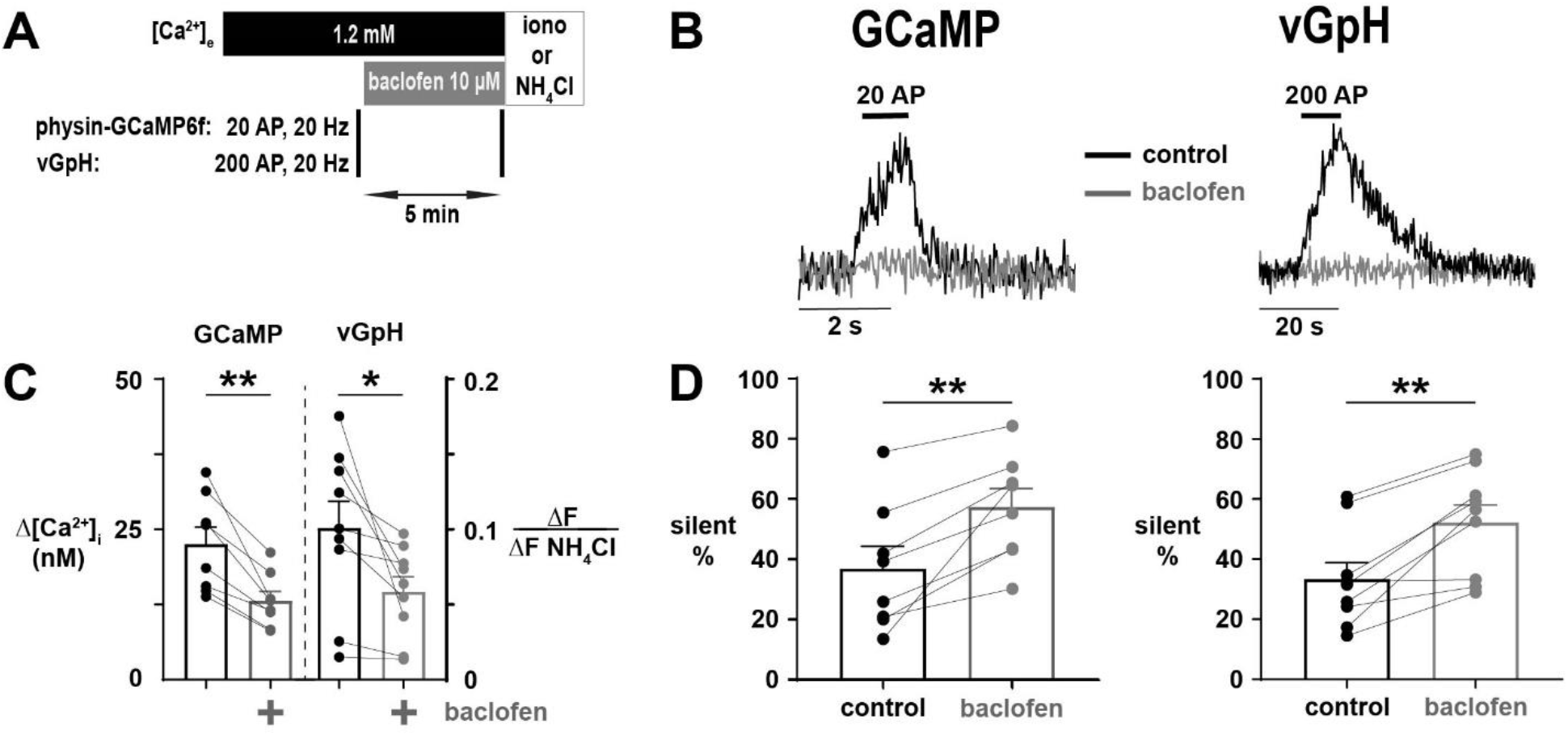
Agonism of GABA_B_R with baclofen leads to silencing of Ca^2+^ influx and SV exocytosis. (A) Diagram of the experimental protocol. (B) Representative traces of silencing of Ca^2+^ influx and SV exocytosis resulting from agonism of GABABR with baclofen in single nerve terminals expressing physin-GCaMP or vGpH, respectively. (C) Δ[Ca^2+^]_i_ and ΔF/ΔF_NH4Cl_ before and following application of baclofen in responding terminals. (D) The percentage of silent terminals before and following treatment with baclofen measured in physin-GCaMP and vGpH-expessing neurons (see labels above). For C and D, dots are individual neurons, bars are mean of all neurons, and error bars are SEM. C and D were analyzed with paired t-test, * p < 0.05, ** p < 0.01, n = 8 for physin-GCaMP and n = 9 for vGpH.

## Discussion

The flux of Ca^2+^ in neurons plays a central role in neurotransmission by triggering the exocytosis of SVs, thereby releasing neurotransmitters into the synaptic cleft^12^. Uncovering the mechanisms regulating Ca^2+^ entry into nerve terminals is therefore critical to understanding the molecular underpinnings of CNS function. The relationship of Ca^2+^ influx and exocytotic rate is highly cooperative^8–11^, highlighting the amplification of neurotransmitter release caused by changes in Δ[Ca^2+^]_i_. Fewer studies have sought to determine the effect of [Ca^2+^]_e_ on Δ[Ca^2+^]_i_ and, to our knowledge, none have examined this in the range encompassing the physiological set point for [Ca^2+^]_e_ at physiological temperature. In general, the narrowing of the AP at warmer temperatures^33,34^ results in lowering of Ca^2+^ influx during AP firing, so additionally working at lower [Ca^2+^]_e_ results in a much lower regime of Δ[Ca^2+^]_i_. Previous attempts (carried out at sub-physiological temperatures) modeled this relationship with Michaelis-Menten kinetics, where Δ[Ca^2+^]_i_ obeyed a simple saturation of VGCCs, but predicted a simple 1:1 relationship as one extrapolates to lower [Ca^2+^]_e_^11,15^.

Here, we show an unexpected but impactful behavior of Ca^2+^ influx in the lower Δ[Ca^2+^]_i_ regime, which is that small modulation of Ca^2+^ influx leads to dramatic silencing of individual nerve terminals. We discovered that a substantial subset of terminals (~40%) exhibited silencing of Ca^2+^ influx and SV exocytosis at physiologic [Ca^2+^]_e_, and this proportion increased steeply (to ~60%) as [Ca^2+^]_e_ was lowered to 0.8 mM. In comparison, the change in the proportion of silent terminals in the transition from [Ca^2+^]_e_ 2.0 mM to 1.2 mM, a difference of ~15%, was smaller despite a larger step in [Ca^2+^]_e_. Previous estimates of synaptic silencing were drawn from experiments conducted at [Ca^2+^]_e_ 2.0 mM and agree with our findings at this concentration^17,22,35,36^. However, our results highlight that the proportion of silent terminals in cultured neurons is substantially higher under conditions of physiologic [Ca^2+^]_e_. Thus, [Ca^2+^]_e_ is a crucial variable to consider in future studies examining synaptic silencing. The agreement of our findings in silencing of both Δ[Ca^2+^]_i_ and SV exocytosis indicates that silencing of Δ[Ca^2+^]_i_ is driving the effect as opposed to an independent mechanism operating on SV exocytosis alone.

The proportion of silencing across different neurons was steeply related to absolute Δ[Ca^2+^]_i_ regardless of VGCC subtypes, such that lowering of Δ[Ca^2+^]_i_ was associated with increasing proportions of silent terminals. Due to this effect, our analysis suggests that the minimal [Ca^2+^]_e_ needed to sustain any neurotransmission is ~ 0.5 mM. Moreover, the degree of silencing caused by acutely blocking VGCC subtypes followed this relationship. Taken together, our results suggest the operation of a feedback mechanism causing the selective shutdown of terminals not meeting a threshold for Ca^2+^ entry. This mechanism has important implications for the impact of neuromodulators that regulate I_Ca_ because relatively small changes in Δ[Ca^2+^]_i_ could cause a substantial proportion of terminals to become silent^28^. As an example, we investigated the impact on synaptic function resulting from inhibition of presynaptic VGCCs by Gβγ subunits using agonism of GABA_B_R by baclofen. Previous studies have demonstrated that the Gβγ interacts with presynaptic VGCCs to lower I_Ca_^31,32,37^. We observed that, in responding terminals, baclofen was associated with ~40% decrease in Δ[Ca^2+^]_i_. This decrement in Ca^2+^ influx led to an increase in the proportion of silent terminals from ~40% to 60%. Notably, this difference in the degree of silencing and Δ[Ca^2+^]_i_ agreed excellently with results from decreasing [Ca^2+^]_e_ 1.2 to 0.8 mM, supporting the conclusion that the effect of baclofen is mediated by decreasing presynaptic Ca^2+^ influx.

While our findings have shown that [Ca^2+^]_e_ contributes importantly to presynaptic neuromodulation, we have not identified the precise molecular mechanism. Previous investigations indicate that silencing is a drastic form of synaptic homeostasis, as chronic suppression of neuronal activity with tetrodotoxin (TTX) decreases the proportion of silent terminals^36^ and increasing activity with chronic depolarization increases this proportion^35,36^. Several intracellular signaling pathways have been identified to set the fraction of silent terminals, including those involving adenylyl cyclase^16^, calcium/calmodulin-dependent protein kinase II^38^, and calcineurin/cyclin-dependent kinase 5^18,22^. The disparate molecular pathways that converge on the regulation of synaptic silencing suggest that it is an important mode of plasticity for neurons in the CNS. Notably, in experiments investigating the impact of GPCR modulation, only a subset of GPCRs tested were able to alter silencing^39^. The silencing caused by GABABR agonism was dependent on proteosome function, and the authors concluded that protein degradation was necessary^39^. However, these experiments utilized 4-hour incubation with baclofen, as opposed to 5 minutes here, so an acute effect of GABA_B_R agonism independent of proteosomal activity may have been missed. Indeed, silencing may have both proteosome-dependent and independent pathways, operating on slow (hours) or acute (< 1 hour) time scales, respectively^40^. Moreover, pharmacologic inhibition of proteosome function prevents depolarization-induced silencing, so the effect of proteosome inhibition may not be specific to GABA_B_R agonism^41^. Thus, our results suggest a separate, rapid mechanism by which nerve terminals can be silenced, adding to the diversity of pathways by which this form of plasticity is achieved.

Our findings highlight that the physiologic set point of [Ca^2+^]_e_ in the CNS functions as an important lever in the modulation of synaptic function because further lowering of Δ[Ca^2+^]_i_ substantially increases the proportion of silent terminals. Although [Ca^2+^] of the CSF is tightly regulated globally^42–44^, models of the synapse that account for its limited volume and restriction of diffusion have predicted that drastic reductions of local [Ca^2+^]_e_ may occur during repetitive presynaptic firing^45,46^. Thus, the phenomenon we observed may be important *in vivo* as a homeostatic mechanism to prevent excessive or runaway synaptic activity^47^. Our results suggest that the magnitude of decrease in [Ca^2+^]_e_ does not need to be large to shut down a sizable proportion of terminals.

In summary, we have utilized highly sensitive, genetically-encoded fluorescent biosensors to dissect the effects of [Ca^2+^]_e_ near the physiologic set point on presynaptic function. We discovered that [Ca^2+^]_e_ has a potent effect in setting the proportion of silent terminals which is driven by Δ[Ca^2+^]_i_. These findings provide evidence that [Ca^2+^]_e_ is an important lever contributing to neuromodulation. Future studies will address the intracellular molecular targets responsible for this acute mechanism of synaptic silencing.

## Materials and Methods

### Animals

All experiments involving animals were performed in accordance with protocols approved by the Weill Cornell Medicine Institutional Animal Care and Use Committee. Neurons were derived from Sprague-Dawley rats (Charles River Laboratories strain code: 001, RRID: RGD_734476) of either sex on postnatal days 0 to 2.

### Neuronal culture

Primary neuronal cultures were prepared as previously described^48^. Hippocampal CA1 to CA3 regions were dissected, dissociated, and plated onto poly-L-ornithine-coated coverslips. Plating media consisted of minimal essential medium, 0.5% glucose, insulin (0.024 g/l), transferrin (0.1 μg/l), GlutaMAX™ 1%, N-21 (2%), and fetal bovine serum (10%). After 1 to 3 days in vitro (DIV), cells were fed and maintained in media with the following modifications: cytosine β-D-arabinofuranoside (4 μM) and FBS 5%. Cultures were incubated at 37°C in a 95% air / 5% CO2 incubator. Calcium phosphate-mediated gene transfer was performed on DIV 6-8, and neurons were used for experiments on DIV 14-21.

### Plasmid constructs

The following published DNA constructs were used: VGLUT1-pHluorin (vGpH)^25^, synaptophysin-GCaMP6f (physin-GCaMP)^49^, and GPI iGluSnFR3 v857(iGluSnFR)^27^, which was a gift from Kasper Podgorski. mRuby-synapsin (Addgene plasmid #187896) was generated by removing GFP from GFP-synapsin^50^ using restriction sites AgeI and BGIII, and substituting it in frame with mRuby obtained from pKanCMV-mRuby3-18aa-actin, which was a gift from Michael Lin (Addgene plasmid #74255).

### Live-cell imaging

Coverslips were loaded onto a custom chamber and perfused at 100 μl min^-1^ via syringe pump (Fusion 4000, Chemyx) with Tyrode’s solution containing (in mM, except if noted otherwise): NaCl 119, KCl 2.5, glucose 30, HEPES 25, CaCl2 0.8-2.0 with MgCl2 adjusted to maintain divalence of 4, D,L-2-amino-5-phosphonovaleric acid (APV) 50 μM, 6-cyano-7-nitroquinoxaline-2,3-dione (CNQX) 10 μM, adjusted to pH 7.4. Temperature was maintained at 37°C with a custom-built objective heater under feedback control (Minco). Fluorescence was stimulated with OBIS 488 nm LX or OBIS 561 nm LS lasers (Coherent) passing through a laser speckle reducer (LSR 3005 at 12° diffusion angle, Optotune). Live-cell imaging was performed with a custom-built, epifluorescence microscope. Emission was acquired with a 40x, 1.3 numerical aperture objective (Fluar, Zeiss) and Andor iXon+ Ultra 897 electron-multiplying charge-coupled device camera camera. APs were evoked with platinum-iridium electrodes generating 1 ms pulses of 10 V cm^-1^ field potentials via a current stimulus isolator (A385, World Precision Instruments). For physin-GCaMP and vGpH, neurons were stimulated with 20 and 200 AP, respectively, delivered at 20 Hz. For iGluSnFR, neurons were stimulated with a single AP, with averaging performed over 15 trials delivered every 20 s. A custom-designed Arduino^®^ board coordinated AP and laser stimulation with frame acquisition. Frame rates for AP recordings with vGpH, GCaMP6f, and iGluSnFR were 5, 50, and 100 Hz, respectively. To achieve frame rates of 50 and 100 Hz, a subregion of the EMCCD chip was used (347 pixels and 169 pixels, respectively, compared to 512 pixels). In experiments with neurons expressing vGpH, Tyrode’s solution with NH4Cl 50 mM replacing an equimolar concentration of NaCl was perfused at the end of the experiment to alkalinize intra-vesicular pHluorin molecules^24^ and enable normalization of fluorescence to the total pool of internal vesicles^18^. Because vGpH has low surface accumulation and fluorescence is therefore largely quenched before AP stimulation^51^, a brief (< 1 min) exposure to NH4Cl was performed before experiments to identify transfected neurons and then washed out for at least 5 min. In experiments with physin-GCaMP, the chamber perfusate was exchanged for Tyrode’s solution with Ca^2+^ 4 mM, pH 6.9, and ionomycin 500 μM to saturate the fluorophore and allow conversion of fluorescence to absolute [Ca^2+^]_i_ ^49^. In experiments with iGluSnFR, glutamate 100 mM was applied to saturate the sensor and normalize to maximum fluorescence^27^. Recordings of physin-GCaMP6f and vGpH included ~200 nerve terminals on average while recordings of iGluSnFR included ~50 nerve terminals due to the limited EMCCD dimensions at this frame rate.

### Key Pharmacological Reagents

Ionomycin, ω-conotoxin GVIA, ω-agatoxin IVA, and SNX-482 were purchased from Alomone labs (catalogue numbers I-700, C-300, STA-500, and RTS-500 respectively). The toxins were applied for three minutes in Tyrode’s solution at concentrations of 1 μM, 400 nM, and 500 nM for ω-conotoxin GVIA, ω-agatoxin IVA, and SNX-482, respectively^19–21^. Baclofen was purchased from Sigma-Aldrich (catalogue number B 5399) and continuously perfused at 10 μM beginning five minutes before AP stimulation^32^.

### Image analysis

Images were analyzed with ImageJ^52^. Circular regions of interest (ROIs) were semi-automatically placed around puncta corresponding to nerve terminals using a custom-written macro. Fluorescence within ROIs was background corrected by subtraction of signal from adjacent, non-synaptic ROIs using a custom-written macro. Images of individual 1 AP trials of iGluSnFr were averaged using a custom-written macro.

### Experimental design and statistical analyses

Quantification of peak fluorescence responses and the percentage of silent terminals was performed with Excel. Terminals were classified as silent if the peak (ΔF) above the baseline (mean of 49 frames) was less than the standard deviation of the baseline (σ_baseline_, 49 frames)^18^. For physin-GCaMP and vGpH, the peak was calculated as the mean of 5 frames at the end of the AP train. For iGluSnFR, the peak was calculated as the mean of 3 frames, with the first frame in the range coinciding with the AP. For vGpH, fluorescence was normalized to the total internal pool of vesicles following perfusion of NH4Cl^18^. For iGluSnFR, fluorescence was normalized to the maximum following perfusion of a saturating concentration of glutamate^27^. For physin-GCaMP, fluorescence was converted to absolute [Ca^2+^]_i_ using^49^:

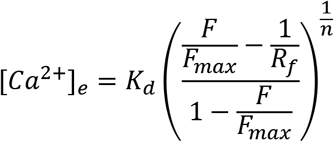

where F_max_ is the peak following saturation of GCaMP6f with ionomycin and K_d_ (0.38 μM), Rf (51.8), and n (2.3) are the *in vitro* dissociation constant, dynamic range, and Hill coefficient, respectively^53^. Statistical analysis was performed with Prism version 8. Comparisons were made with t-test or, for more than two groups, ANOVA with Tukey’s post-test for multiple comparisons. Peak responses of iGluSnFR in silent terminals were compared with a mixed-effects model because some neurons lacked silent terminals at [Ca^2+^]_e_ 2.0 mM (Figure 6E). Statistical significance was defined as P < 0.05. The relationship of Δ[Ca^2+^]_i_ in responding terminals to the proportion of silent terminals was fit with a Hill equation using least squares regression and constraining the maximum silencing at Δ[Ca^2+^]_i_ = 0 to 100% (Figures 4E and F). Data are presented as mean ± SEM except in Figure 1D, which demonstrates mean ± 95% confidence interval to illustrate the experimental difference from the theoretical value.

## Acknowledgements

We thank members of the Ryan lab for their valuable discussion of the manuscript. We give special thanks to Andrew Nelson for developing the custom Arduino^®^ board and its graphical user interface. We thank Kasper Podgorski for kindly providing iGluSnFR3 v857.

